# ASlive: a database for alternative splicing atlas in livestock animals

**DOI:** 10.1101/769273

**Authors:** Jinding Liu, Suxu Tan, Shuiqing Huang, Wen Huang

## Abstract

We present in this study the development and implementation of a database for alternative splicing atlas in livestock animals (ASlive.org). Alternative splicing is an important biological process whose precision must be tightly regulated during growth and development. Using publicly available RNASeq data sets across many tissues, cell types, and biological conditions totaling 28.6 tera bases, we built a database of alternative splicing events in five major livestock animal species (cattle, sheep, pigs, horses, and chickens). The database contains many types of information on alternative splicing events, including basic information such as genomic locations, genes, and event types, quantitative measurements of alternative splicing in the form of percent spliced in (PSI), overlap with known DNA variants, as well as orthologous events across different lineage groups. This database, the first of its kind in livestock animals, will provide a useful exploratory tool to assist functional annotation of animal genomes.

## Introduction

Splicing of multi-exonic precursor messenger RNAs (pre-mRNA) is a key biological process that can impact both the sequences and expression of proteins. In particular, multi-exonic pre-mRNAs have the potential to be alternatively spliced. Alternative splicing allows one gene to code for multiple mature mRNA and protein isoforms, greatly expanding the diversity of the proteome (Nilsen and Graveley 2010). For example, the Drosophila Down syndrome cell adhesion molecule (*Dscam*) gene is able to generate more than 38,000 possible isoforms with variable immunoglobin and transmembrane domains (Schmucker *et al.* 2000). This remarkable diversity of a transmembrane receptor gene provides the specificity for neuronal connectivity needed in axon guidance. The precise regulation of alternative splicing is important in development and growth. Thus, the disruption of normal alternative splicing can lead to diseases such as cancers. Indeed, natural DNA variation that results in genetic variation in alternative splicing is a major determinant of phenotypic diversity among individuals in a population, including genetic risks to diseases (Li *et al.* 2016). In livestock animals, where genetic improvement is a major goal, the specific role of alternative splicing in determining phenotypic variation in economic traits is not well understood. Part of the reason is the lack of a comprehensive annotation of alternative splicing in these agricultural species. For example, while the size of the genome (3.1 Gbp for humans and 2.7 Gbp for cattle) and number of protein coding genes (20,454 for humans and 21,880 for cattle) are similar for humans and cattle, there are on average 5.1 annotated splice isoforms per human gene versus 1.6 per cattle gene, a more than three-fold difference (Zerbino *et al.* 2018).

The advent of high throughput sequencing technologies has greatly facilitated genome annotation efforts. In addition, targeted experimental studies have increasingly utilized next generation sequencing to globally survey the transcriptomes of different cell types, tissues, and animals across many organisms. Such diversity of experimental data provides unprecedented breadth and depth of transcriptomes across many species in public databases, including livestock animals. However, most studies focus on differences in steady state RNA abundance, which represents an equilibrium between transcription and mRNA decay and does not capture difference in post-transcriptional regulation such as splicing.

Experimental data in public databases such as the sequence read archive (SRA) are highly heterogeneous. While this presents a challenge to re-use these data, it also provides a great opportunity to discover new information, some of which only happens in specific conditions. As such, heterogeneous and diverse experimental data in public databases complement organized annotation projects that typically only use limited samples and conditions. For example, even for humans, experimental data in the SRA database contained a large number of unannotated splice junctions (Nellore *et al.* 2016).

There are several alternative splicing specific databases available. For instance, the ASpedia (Alternative Splicing Encyclopedia of Human) database contains a collection of alternative splicing events identified from a single project with 26 tissues and 241 samples (Hyung *et al.* 2018). The CancerSplicingQTL is a database to search and browse splicing quantitative trait loci (sQTLs) affecting alternative splicing in cancer samples (Tian *et al.* 2019). These databases become increasingly useful as an exploratory and hypothesis generating tool. However, no database is specifically designed for livestock animals.

In this study, we present the development of the alternative splicing in livestock animals (ASlive) and a web interface for users to interact with the database. There are several unique features of the database. We developed a uniform processing pipeline to process over 4,000 samples in the SRA database, covering 188 tissues in five major livestock animal species (cattle, sheep, pigs, horses, and chicken), totaling 28.6 tera bases of sequence data. We discovered hundreds of thousands of unannotated alternative splicing events that were supported by multiple lines of experimental evidence and quantitatively estimated their alternative splicing level. We also identified conservative alternative splicing events across species, allowing users to assess and explore the tissue and species specificity of alternative splicing events. This study provides an important new tool to the animal genome research community and complements ongoing large-scale annotation projects such as the functional annotation of animal genomes (FAANG) project (Andersson *et al.* 2015).

## Data collection and processing

### Data Collection

The reference genome assemblies of five livestock species including cattle (taxonomy id: 9913), sheep (9940), pigs (9823), horses (9796) and chicken (9031) were downloaded from Ensembl (release 96). We also obtained reference annotations from both Ensembl and RefSeq. Sequence data from a total of 4,166 RNASeq experiments containing 8,257 runs and 28.6 tera bases in the SRA database were collected by querying the meta data of the SRA database (Table 1). To simplify our data processing pipeline, we restricted data to the Illumina platform, which constituted the vast majority of RNASeq data.

**Table 1.**
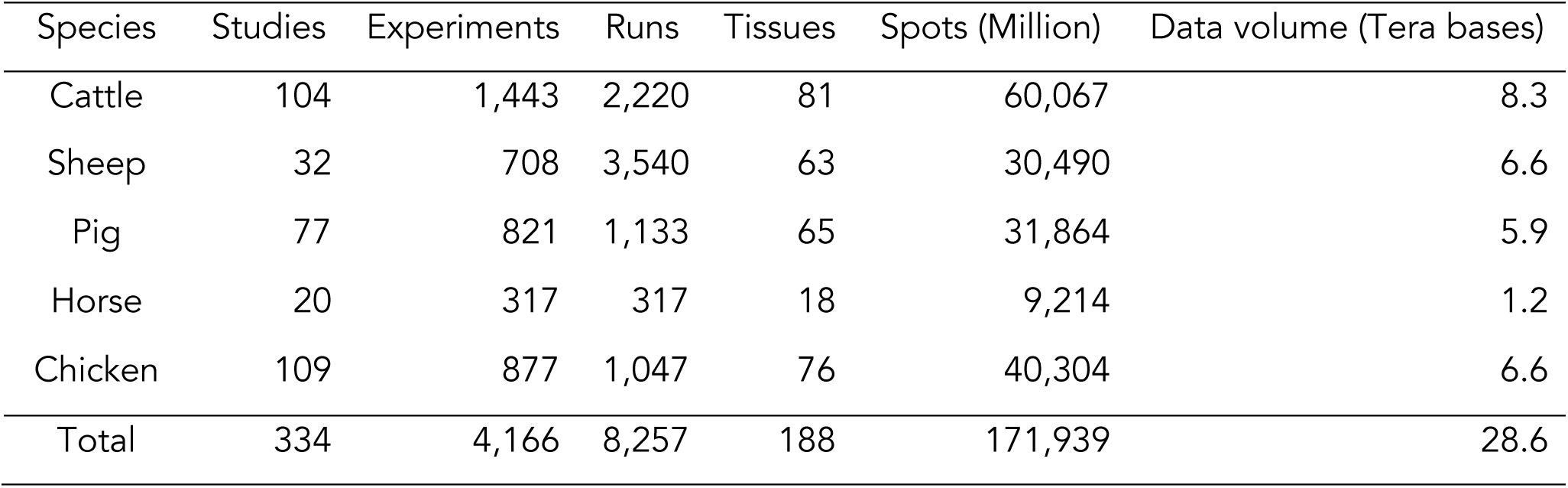
Summary of RNASeq data used in ASlive

### Improvement of gene models

The reference annotations from Ensembl and RefSeq were largely incomplete for livestock species. We used the following procedure to improve the annotations using high quality RNASeq data from SRA (Table 2).

**Table 2.** Summary of improvement of gene models.

| Species | Ensembl+RefSeq |  |  | SRA data used |  | After improvement |  |  |
| --- | --- | --- | --- | --- | --- | --- | --- | --- |
|  | Genes | Transcripts | Transcripts per gene | Total sequenced fragments (M) |  | Genes | Transcripts | Transcripts per gene |
| Cattle | 32,731 | 95,018 | 2.9 | 17,444 |  | 35,661 | 175,198 | 4.9 |
| Sheep | 27,829 | 44,398 | 1.6 | 10,212 |  | 28,974 | 65,191 | 2.2 |
| Pig | 30,284 | 101,216 | 3.3 | 8,982 |  | 31,959 | 157,045 | 4.9 |
| Horse | 35,886 | 111,890 | 3.1 | 2,606 |  | 36,310 | 124,270 | 3.4 |
| Chicken | 27,251 | 81,909 | 3.0 | 16,183 |  | 29,091 | 156,429 | 5.4 |

1. Ensembl and RefSeq annotations were compared using cuffcompare by setting Ensembl as the reference. RefSeq transcripts that were flagged as “j” (novel isoform) and “u” (novel transcribed region) were added to the Ensembl annotation. This merged annotation served as the reference annotation in subsequent steps.
2. Experiments with at least 40 million spots (30 million for horses due to low number of experiments passing the filter) and 75 bp read length were mapped to the reference genome using HISAT2 (Kim *et al.* 2019) in the presence of the reference annotation. Those with at least 40 million mapped fragments were retained and assembled into reference guided gene models in GTF format using StringTie (Pertea *et al.* 2015).
3. We then improved the reference annotation by iteratively comparing each assembled GTF file to the annotation from the previous iteration. Briefly, one assembled GTF file was compared with the GTF file from the previous iteration using cuffcompare. Novel multi-exonic transcripts (“j” and “u”) that were at least 200 bp long, with an average coverage of 2x per transcript, and an average coverage of 1x per exon for all exons were added. This process was iteratively performed through all StringTie assembled GTFs from the previous step.
4. The final filtering step consisted of comparing all GTF files from step 2) to the merged GTF file from step 3) and requiring that all novel transcripts must occur in at least three different studies and four different experiments.

### Identification and quantification of alternative splicing events

After aligning RNASeq reads to the improved reference annotation in each species using HISAT2, we used rMATs (Shen *et al.* 2014) to identify and quantify alternative splicing events in all samples. rMATs reports junction read counts, effective junction length for each alternative splicing event and classifies them into five classes including alternative 5’ splice site (A5SS), skipped exon (SE), mutually exclusive exons (MXE), retained intron (RI), and alternative 3’ splice site (A3SS). It is important to note that rMATs is highly sensitive and does not rely on the GTF annotation to identify alternative splicing events and may report events that do not conform to existing intron chains in the annotation. We retained these events in our database because they were supported by junction reads. Alternative splicing events from all samples were merged to create a non-redundant catalog. To further refine the catalog, we retained events that were evident by at least three skipping reads and three inclusion reads in at least four different experiments and three different studies (Table 3). We identified between 48,208 and 151,087 confident alternative splicing events in each of the five species (Table 3). Quantitative measurements including the percent spliced in (PSI), numbers of skipping and inclusion reads, and the effective junction lengths were collected.

**Table 3.** Summary of alternative splicing events identified from SRA data

| Species | A5SS | SE | MXE | RI | A3SS | Total |
| --- | --- | --- | --- | --- | --- | --- |
| Cattle | 10,227 | 82,153 | 25,130 | 20,364 | 13,213 | 151,087 |
| Sheep | 1,567 | 50,030 | 11,148 | 2,449 | 2,390 | 67,584 |
| Pig | 8,652 | 68,309 | 23,876 | 17,723 | 11,107 | 129,667 |
| Horse | 3,176 | 29,564 | 6,164 | 4,358 | 4,946 | 48,208 |
| Chicken | 10,088 | 58,752 | 19,415 | 19,892 | 12,128 | 120,275 |

### Identification of orthologous alternative splicing events

To enable comparative analyses, we first identified alternative splicing events that are orthologous among the livestock species. All alternative splicing events including those without sufficient experimental support were considered in this step because they may have support based on orthology. We lifted coordinates of exon boundaries over to the human genome assembly (hg38) using the LiftOver tool from UCSC Genome Browser (Haeussler *et al.* 2019) for all species. This allowed us to use the hg38 coordinate system as a reference to identify 1:1:1:1:1 orthologous exons across all five species, *i.e.*, there were unique reciprocal alignments of exons. To identify orthologous alternative splicing events, we searched the coordinates of the intron chains across groups of species. An alternative splicing event was considered orthologous among a group if it was present in all species in the group. We considered orthology at four phylogenetic levels, including 17,639 orthologous events in bovida (cattle and sheep), 8,961 in artiodactyla (cattle, sheep and pigs), 5,352 in mammals (cattle, sheep, pigs, and horses), and 3,276 in vertebrates (all fives species) (Table 4). The most abundant type of conservative alternative splicing events is the skipped exon (SEs).

**Table 4.** Summary of conservative alternative splicing events

| Lineage | A5SS | SE | MXE | RI | A3SS | Total |
| --- | --- | --- | --- | --- | --- | --- |
| Vertebrate | 21 | 3,126 | 97 | 0 | 32 | 3,276 |
| Mammal | 42 | 4,927 | 272 | 9 | 102 | 5,352 |
| Artiodactyla | 22 | 8,038 | 840 | 6 | 55 | 8,961 |
| Bovidae | 85 | 14,660 | 2,606 | 51 | 237 | 17,639 |

### Web interface

A simple and intuitive web interface (ASlive.org) was designed for users to explore the ASlive database (Figure 1a). There are two primary ways to initiate a query against the database, which are easily accessible within a navigation bar of the ASlive website (Figure 1a). Users may search the database by entering the specific genomic locations, gene symbols, or Pfam and GO annotations (Figure 1b). Alternatively, the database can be queried by blasting a sequence (Figure 1c). This is particularly useful when looking for orthologous genes in a different species when they are not easily identified by gene symbols. Both entry points lead to similarly structured list of alternative splicing events that match the query. The results of the search are displayed in a concise table form (Figure 2a). The table (Figure 2a) can be downloaded for further analyses by the users. Within the table, users may refine the research results by imposing additional search criteria, open a pop-up window to explore the details of the alternative splicing events, and link to the Ensembl genome browser.

**Figure 1.**
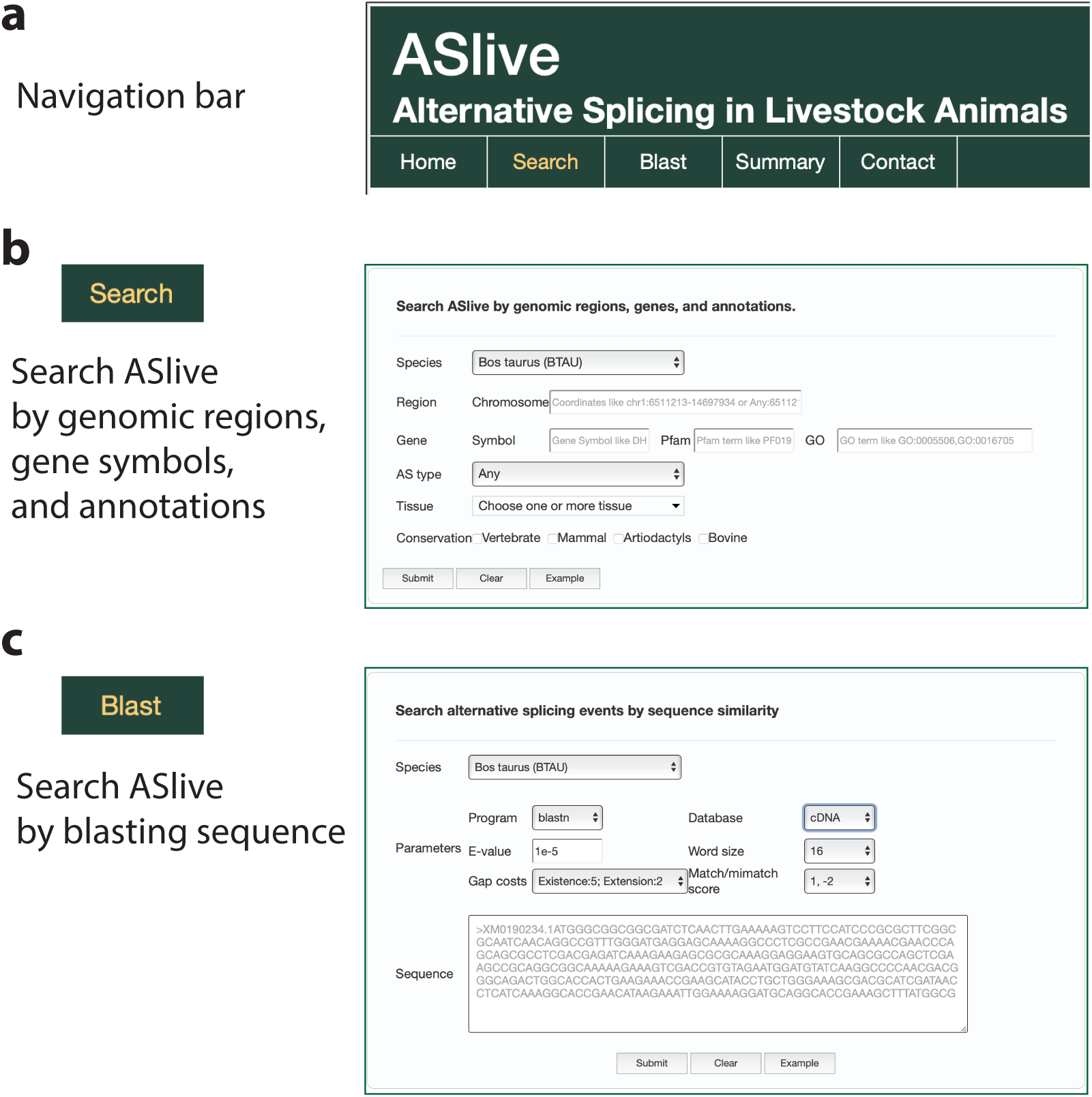
Web interface of ASlive. **(a)** Navigation bar of the web interface for ASlive.org. **(b)** Entry point for the database by search based on genomic locations, gene symbols, and annotations. **(c)** Entry point for the database by search based on sequence similarity.

**Figure 2.**
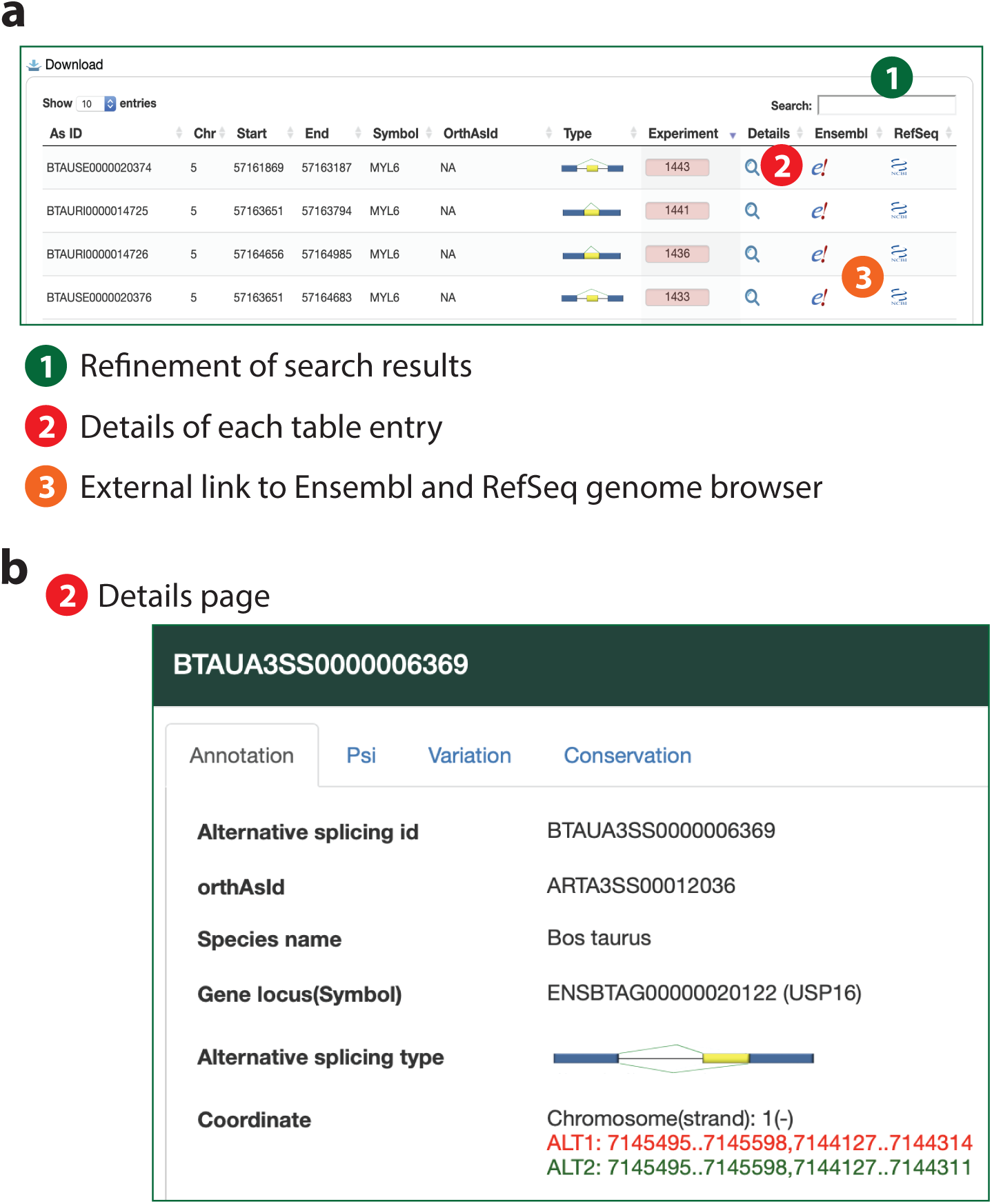
Information and data contained in ASlive. **(a)** Display of search results and links to additional information in ASlive. **(b)** Basic information on alternative splicing events and tabs in the details page that leads to additional information including PSI, overlap with DNA variation, and conservation.

The details window for each alternative splicing event contains a wealth of information we gathered from either the SRA data or other databases. There are four tabs in this window. First, the annotation tab provides basic information for the event including unique ID, orthologous ID if available, classification of the event and the coordinates of exon boundaries that are involved in the splicing (Figure 2b). Second, the Psi tab offers the PSI data across all SRA experiments with tissue annotation in a sortable table. A box plot showing the variation across experiments and tissues is also displayed (Figure 3a). Third, the variation tab provides a list of dbSNP variants that overlap within the exons and introns of the alternative splicing event, including whether they overlap with the acceptor/donor sites. Finally, the conservation tab provides a boxplot visualization of PSIs across species where the event is conserved (Figure 3b). These data visualizations allow users to quickly assess the biological significance of an alternatives splicing event, such as whether it is conserved or specific across tissues and species. Users may also download data associated with these visualizations to explore further details.

**Figure 3.**
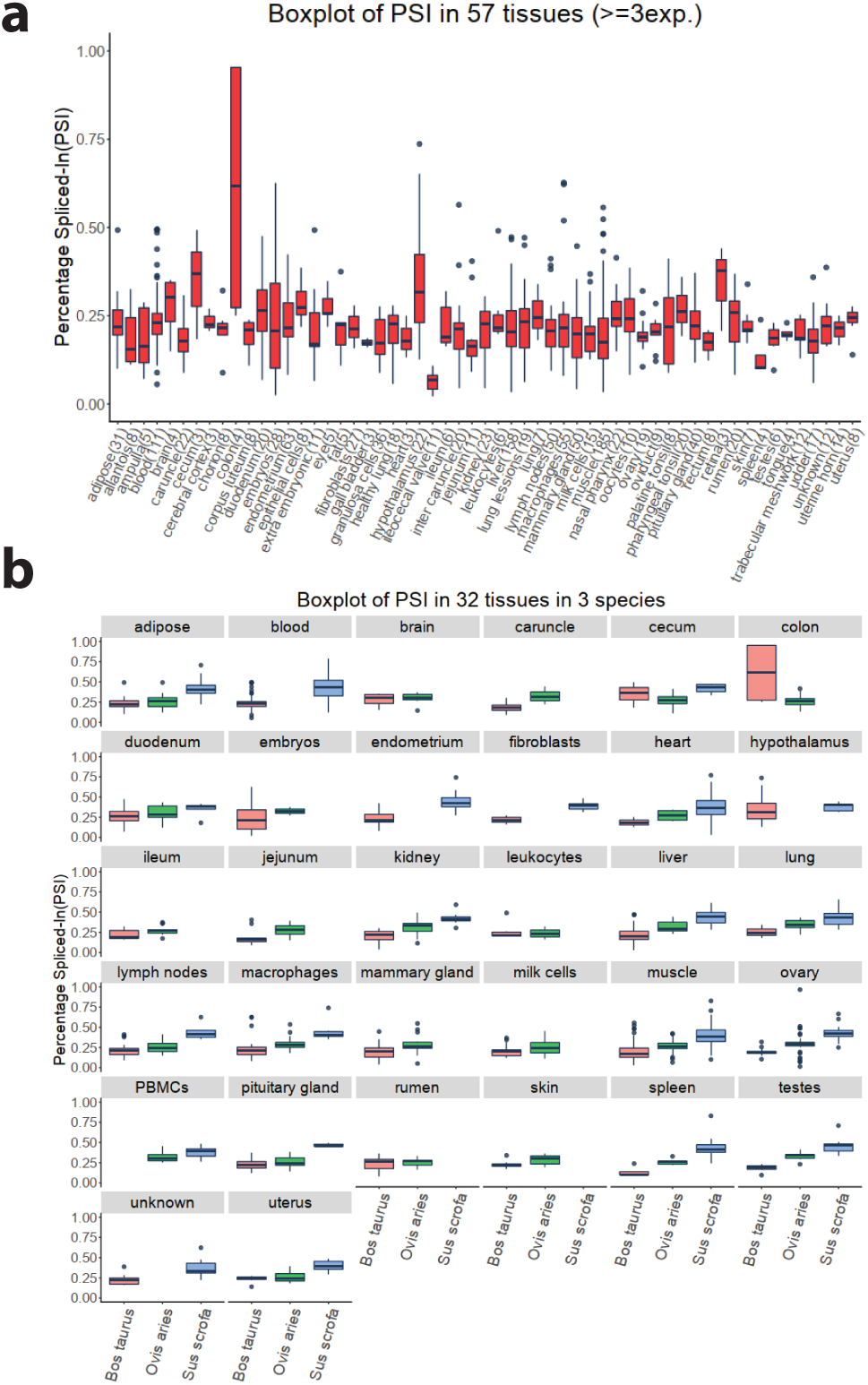
Visualization of quantitative alternative splicing information across tissues and species. Boxplots are used to display the variation within and across 57 tissues of an alternative splicing event in bovine **(a)** and the same information in 32 tissues in three species for the same event **(b).**

## Discussion

We describe the development and implementation of a comprehensive alternative splicing database in livestock animals - ASlive.org. The database fills an important gap in the current literature and web space and has several unique features. For example, it is the first database specifically designed for livestock animals to capture alternative splicing events in heterogeneous samples, which allows users to obtain experimental support of alternative splicing events from a wide range of tissues, cell types, and biological conditions. Unlike many other alternatives splicing databases which relies on a good assembly (typically in GTF format) to identify alternative splicing events, we used rMATs to also identify novel events that is independent of transcript assemblies. Second, we design the interface to meet various needs, including experimental biologists who focus on the details of a small number of genes or computational scientists who are interested in downloading the primary data and processing them offline. Third, we present one of the first databases to include orthologous alternative splicing events, which cannot be easily accessed through existing genome browsers and databases.

As RNASeq data in data archives grow, we plan to regularly update the database with new data. Our ID system of alternative splicing events allows us to add new events without altering existing IDs, providing backward compatibility. Nevertheless, the existing data already have a comprehensive coverage of tissues, cell types, and biological conditions and likely will serve most purposes. Because of the important role of genetic variation in animal related research, we plan to incorporate additional data sources that can capture the relationship among genetic variation at the DNA, splicing, and phenotypic levels. This could be, for example, achieved by incorporating genotype-phenotype associations present in the animal QTLdb (https://www.animalgenome.org) (Hu *et al.* 2019).

## Acknowledgement

Funding for this study was provided by Michigan State University AgBioResearch (W.H.) and Nanjing Agricultural University (KYZ201667, J.L.).

## References

Andersson L., A. L. Archibald, C. D. Bottema, R. Brauning, S. C. Burgess, et al., 2015 Coordinated international action to accelerate genome-to-phenome with FAANG, the Functional Annotation of Animal Genomes project. Genome Biol. 16: 57. https://doi.org/10.1186/s13059-015-0622-4

Haeussler M., A. S. Zweig, C. Tyner, M. L. Speir, K. R. Rosenbloom, et al., 2019 The UCSC Genome Browser database: 2019 update. Nucleic Acids Res. 47: D853–D858. https://doi.org/10.1093/nar/gky1095

Hu Z.-L., C. A. Park, and J. M. Reecy, 2019 Building a livestock genetic and genomic information knowledgebase through integrative developments of Animal QTLdb and CorrDB. Nucleic Acids Res. 47: D701–D710. https://doi.org/10.1093/nar/gky1084

Hyung D., J. Kim, S. Y. Cho, and C. Park, 2018 ASpedia: a comprehensive encyclopedia of human alternative splicing. Nucleic Acids Res. 46: D58–D63. https://doi.org/10.1093/nar/gkx1014

Kim D., J. M. Paggi, C. Park, C. Bennett, and S. L. Salzberg, 2019 Graph-based genome alignment and genotyping with HISAT2 and HISAT-genotype. Nat. Biotechnol. 37: 907–915. https://doi.org/10.1038/s41587-019-0201-4

Li Y. I., B. van de Geijn, A. Raj, D. A. Knowles, A. A. Petti, et al., 2016 RNA splicing is a primary link between genetic variation and disease. Science (80-.). 352: 600–604. https://doi.org/10.1126/science.aad9417

Nellore A., A. E. Jaffe, J.-P. Fortin, J. Alquicira-Hernández, L. Collado-Torres, et al., 2016 Human splicing diversity and the extent of unannotated splice junctions across human RNA-seq samples on the Sequence Read Archive. Genome Biol. 17: 266. https://doi.org/10.1186/s13059-016-1118-6

Nilsen T. W., and B. R. Graveley, 2010 Expansion of the eukaryotic proteome by alternative splicing. Nature 463: 457–463. https://doi.org/10.1038/nature08909

Pertea M., G. M. Pertea, C. M. Antonescu, T.-C. Chang, J. T. Mendell, et al., 2015 StringTie enables improved reconstruction of a transcriptome from RNA-seq reads. Nat Biotechnol. 33: 290–295. https://doi.org/10.1038/nbt.3122

Schmucker D., J. C. Clemens, H. Shu, C. A. Worby, J. Xiao, et al., 2000 Drosophila Dscam is an axon guidance receptor exhibiting extraordinary molecular diversity. Cell 101: 671–84.

Shen S., J. W. Park, Z. Lu, L. Lin, M. D. Henry, et al., 2014 rMATS: Robust and flexible detection of differential alternative splicing from replicate RNA-Seq data. Proc. Natl. Acad. Sci. 111: E5593–E5601. https://doi.org/10.1073/pnas.1419161111

Tian J., Z. Wang, S. Mei, N. Yang, Y. Yang, et al., 2019 CancerSplicingQTL: a database for genome-wide identification of splicing QTLs in human cancer. Nucleic Acids Res. 47: D909–D916. https://doi.org/10.1093/nar/gky954

Zerbino D. R., P. Achuthan, W. Akanni, M. R. Amode, D. Barrell, et al., 2018 Ensembl 2018. Nucleic Acids Res. 46: D754–D761. https://doi.org/10.1093/nar/gkx1098

